# Cellular structure image classification with small targeted training samples

**DOI:** 10.1101/544130

**Authors:** Dali Wang, Zheng Lu, Yichi Xu, Zi Wang, Chengcheng Li, Anthony Santella, Zhirong Bao

## Abstract

**Motivation:** Cell shapes provide crucial biology information on complex tissues. Different cell types often have distinct cell shapes, and collective shape changes usually indicate morphogenetic events and mechanisms. The identification and detection of collective cell shape changes in an extensive collection of 3D time-lapse images of complex tissues is an important step in assaying such mechanisms but is a tedious and time-consuming task. Machine learning provides new opportunities to automatically detect cell shape changes. However, it is challenging to generate sufficient training samples for pattern identification through deep learning because of a limited amount of images and annotations.

**Result:** We present a deep learning approach with minimal well-annotated training samples and apply it to identify multicellular rosettes from 3D live images of the *Caenorhabditis elegans* embryo with fluorescently labelled cell membranes. Our strategy is to combine two approaches, namely, feature transfer and generative adversarial networks (GANs), to boost image classification with small training samples. Specifically, we use a GAN framework and conduct an unsupervised training to capture the general characteristics of cell membrane images with 11,250 unlabelled images. We then transfer the structure of the GAN discriminator into a new Alex-style neural network for further learning with several dozen labelled samples. Our experiments showed that with 10-15 well-labelled rosette images and 30-40 randomly selected non-rosette images our approach can identify rosettes with over 80% accuracy and capture over 90% of the model accuracy achieved with a training dataset that is five times larger. We also established a public benchmark dataset for rosette detection. This GAN-based transfer approach can be applied to study other cellular structures with minimal training samples.

## 1 Introduction

Live microscopy and image processing are commonly used to investigate cellular dynamics, quantify cellular behaviors, and support simulation-based hypothesis testing. The huge amount of microscope data generated during the studies presents unprecedented challenges for human-based, interactive data analysis. Advanced computing technology has been used in microscopic data analysis (Jones *et al*., 2009); however, the majority of these efforts require deep domain knowledge through a label-intensive annotation process. Nowadays, AI-based computer vision provides a “model-free” approach to solving generic data problems, such as object identification. For example, convolutional neural networks (CNNs) are widely adopted for object classification and identification (Krizhevsky *et al*., 2012; Szegedy *et al*., 2015; Simonyan and Zisserman, 2014). Some well-known CNNs usually contain a large number of parameters (e.g., more than 25 million in a ResNet-50 network), which require large well-labelled training datasets. However, considering funding limitations and the scarcity of domain experts, it is still quite challenging to establish comprehensive training datasets from 3D live images of complex tissues for detecting particular cellular structures.

Multicellular rosette is a form of collective cell shape change that is used to mediate tissue morphogenesis in diverse organisms and biological processes, such as cell intercalation, collective migration, and collective outgrowth of neurites (Blankenship *et al*., 2006; Harding *et al*., 2014; Fan *et al*., 2018). Multicellular rosettes form when neighboring cells contract their contact so that multiple cells converge at the center (Fig. 1). Because these structures are transient and rare, automated detection is essential to identifying them and characterizing their dynamics (Farrell *et al*., 2017).

**Fig. 1.**
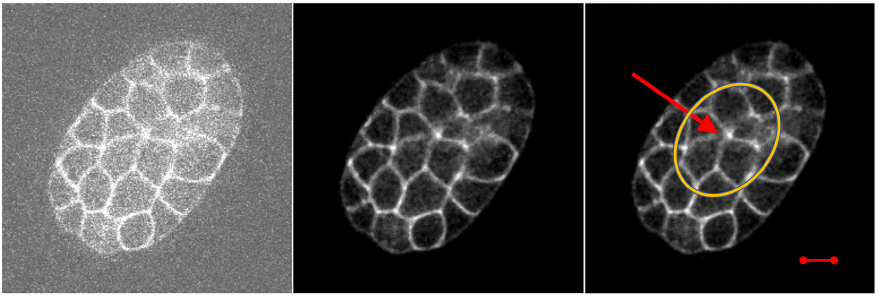
Microscopy image of C. elegans before and after denoising. The orange circle in the left image contains a rosette structure formed by five adjacent cells with a common center (marked by a red arrow). The short red line at the right bottom of the image represents a 32-pixel segment.

We present a method for unique cellular structure image classification using 3D time-lapse datasets directly. Our learning process consists of two steps: common cellular structure learning with unlabeled datasets (relatively easy to obtain) and target cellular structure learning with small labelled datasets. We adopt basic concepts within the generative adversarial networks (Goodfellow *et al*., 2014; Arjovsky and Bottou, 2017; Odena *et al*., 2016; Li *et al*., 2018; Radford *et al*., 2015) to capture the common structure learning with unlabeled dataset. Then we ingest a small quantity of target structure images into a hybrid classifier to improve the efficiency of target structure learning via transfer learning and regularization (Pan *et al*., 2010; Noroozi and Favaro, 2016; Doersch *et al*., 2015). We also quantify the performance of the hybrid microstructure classifier with different sizes of training samples.

## 2 Methods

### 2.1 Dataset

#### 2.1.1 Raw data

We use a set of *C*. *elegans* microscopic images that contains 45 embryos with ubiquitous fluorescent labeling of cell membrane. The raw images are 512 x 512 pixels in size and may contain one to three embryos. The raw images are arranged in sets, each of which contains 300 image stacks taken at 75-second intervals over the first 375 minutes of embryogenesis. Each stack is a pseudo 3D image that contains 30 slices at 1 um vertical distance covering the entire embryo. Image acquisition followed published protocols (Shah *et al*., 2017).

#### 2.1.2 Image sets for experiments

Because each raw image may contain more than one embryo, we first crop the raw images to 128 x 128 images so that each 128 x 128 image contains the complex global structure information of a single embryo. Technically, we write an ImageJ (Schneider *et al*., 2012) macro for this task: For each embryo, we first mark its bounding box, and then inside this bounding box, we randomly select 128 x 128 images. For each of these images, we apply a 3-D median filter and adjust the brightness range to remove the image noise. Examples of a raw image and denoised image are shown in Fig. 1.

We select the raw data from a early developmental period of 61 to 110 minutes. The embryo structure is relative simple before 61 minutes and a meaningful structural pattern of a rosette is seldom observed. For each image stack, we use images between slice 9 and slice 13 as these slices usually have the best imaging quality.

Our goal is to identify and locate a rosette in the 128 x 128 3D microscopic images. Considering the average cell size during the above-mentioned developmental period and the typical structure of a muticelluar rosette, we determine 32 x 32 to be the appropriate image size for classification and detection.

We create two image datasets for deep learning experiments. The first dataset is a collection of unlabelled images for a unsupervised image synthesis task. The second dataset is a manually labelled dataset for a supervised image classification task for target cellular structures. For the unlabelled dataset, we randomly sample one 32 x 32 image at each slice of the image stack. In total, our unlabelled dataset contains 50-minute live image stacks of 45 *C*. *elegans* embryos. Because five slices of each image stack are collected, there are 45 x 50 x 5 = 11250 image patches in our dataset. The labelled dataset contains 78 manually selected 32 x 32 rosette images from 45 different embryos. Each of the manually annotated rosette images contains a multicelluar convergence center. Around 200 non-rosette images are also randomly selected from the unlabelled datasets (see Section 3.2.1 for more information).

### 2.2 Neural network configuration

We modify an AlexNet-styled convolutional neural network (CNN) (Krizhevsky *et al*., 2012) to classify the image (illustrated in Fig. 2). CNNs use convolutional filters to automatically capture features rather than using hand-engineered features in traditional machine learning algorithms. Because the input of our network is a 32 x 32 grayscale image, our network has three convolutional layers, followed by two fully connected layers. We use 4 x 4 filters for all convolutional layers. The number of filters at the first convolutional layer is 32 and doubled at each convolutional layer. Unlike AlexNet, we replace all pooling layers with stride (2 pixels) convolutions so that the network can learn its own pooling method (Springenberg *et al*., 2014). We also place a batch normalization layer after each convolutional layer and the first fully connected layer. Leaky ReLU is used as the activation function for all layers except the last fully connected layer in the network. This architecture is used for all the image classifiers in our study.

**Fig. 2.**
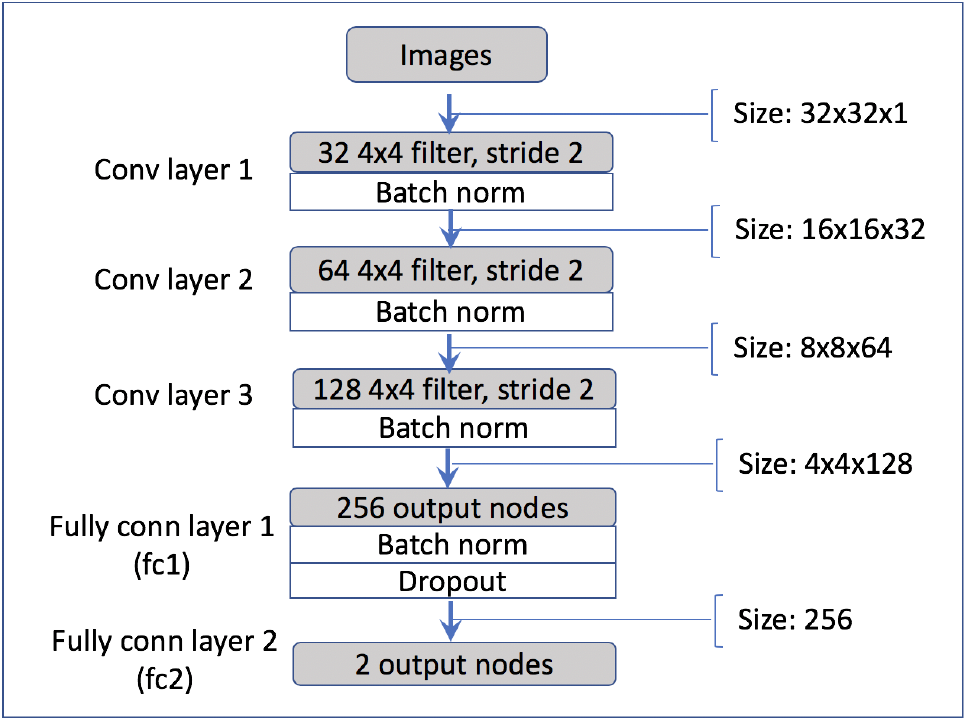
Neural network structure of image classifier.

### 2.3 Generative adversarial networks

We present a way to use a sizable unlabelled dataset and transfer learning techniques to improve the training of the CNNs. Some efforts use pretrained networks and fine-tune them with the small labelled dataset. However, most publicly available networks are pretrained on benchmark computer vision datasets, such as ImageNet (Deng *et al*., 2009). Due to the significant differences between the 3D live images and the images in benchmark datasets, features learned from computer vision datasets are not directly suitable for our scientific dataset. Furthermore, there is no sizable labelled dataset similar in structure to our dataset that can be used to pretrain the network. Hence we take a different approach that uses generative adversarial networks (GANs) and the sizable unlabelled dataset to first learn common features of these images and then transfer the learned features to new CNNs, which are then further tuned with a small labelled dataset.

Generative adversarial networks (GANs) (Goodfellow *et al*., 2014) is a generative framework that consists of two competing networks: a generator network and a discriminator network. The generator produces synthetic data to fool the discriminator, while the discriminator discriminates between real data and synthetic data. The game between the generator *G* and the discriminator *D* is the minimax objective. We use a particular form of GAN, called Wasserstein GAN. A three-convolutional-layer alex-style network structure is used for both the generator and the discriminator. The discriminator network has the same network structure as our classifier shown in Fig. 2.

Samples of generated images patches are shown in Fig. 3(a). These patches are compared with image patches in the real dataset in Fig. 3(b). As shown, the newly generated images in Fig. 3(a) captured the majority of the common features of these live images. The Wasserstein losses for both the generator and discriminator of the 32 x 32 image case are also shown in Fig. 3(c) and 3(d).

**Fig. 3.**
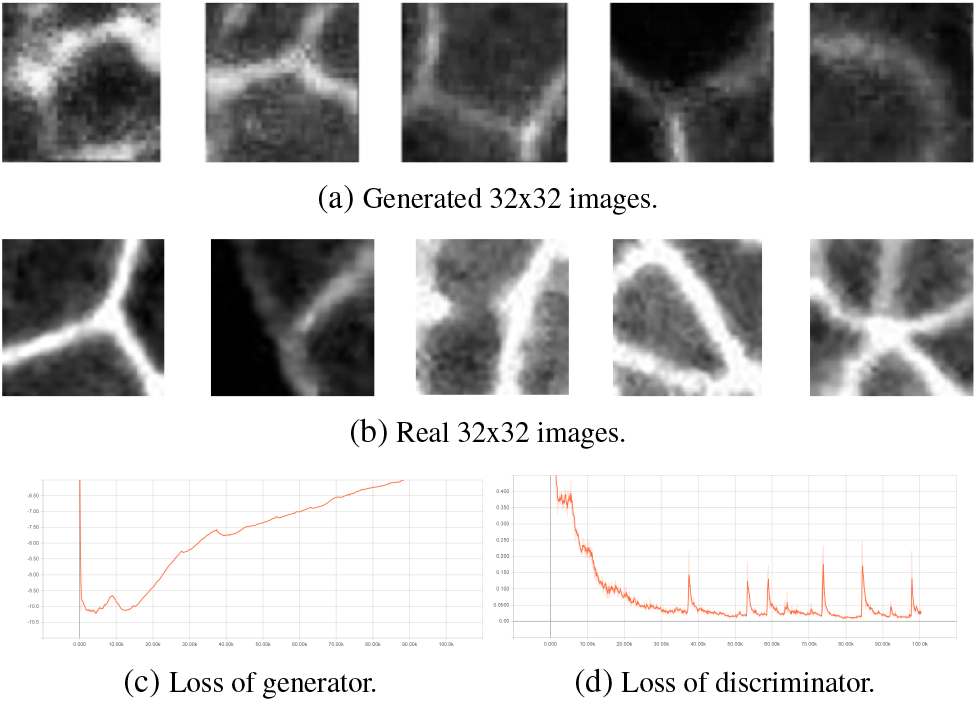
Generated image patches compared with real image patches and associated Wasserstein losses.

### 2.4 GAN-based feature transfer

We first train the GANs with the unlabelled dataset so that the networks can distinguish fake images from real images by learning most of the common features of the unlabelled real images. We then transfer these features to new CNNs and continue to train the new CNNs with labelled images to capture the target cell structures.

Technically, we are interested in the features learned by the GAN discriminator. We create new neural networks, using the same network structure of the discriminator without the last fully connected layer, to capture the learned features from the GAN discriminator. The output of the last convolutional layer contains all the structural features learned by a network. The fully connected layers contain most of the weights in the architecture and a large number of parameters that contain useful information for the target task.

We remove the last layer in the GAN discriminator, which is designed to differentiate the difference between real image and generated images. We then add a new fully connected layer to classify an image with or without a rosette. We use the weights of each convolutional layer and the first fully connected layer of the GAN discriminator (e.g., the fc1 layer in Fig. 2) to initialize the classifier. Because both the classifier and the GAN discriminator use batch normalization after each convolutional layer and the first fully connected layer, there is no bias term for these layers as a new feature of the “layers(contrib)” package provided by tensorflow (Abadi *et al*., 2015). Therefore instead of transferring a bias term of each layer, we transfer parameters in batch normalization layers.

After initializing the neural network with parameters from the GAN discriminator, we continue to train the network using a small manually labelled dataset. The overall workflow of the GAN-based classification is illustrated in Fig. 4.

**Fig. 4.**
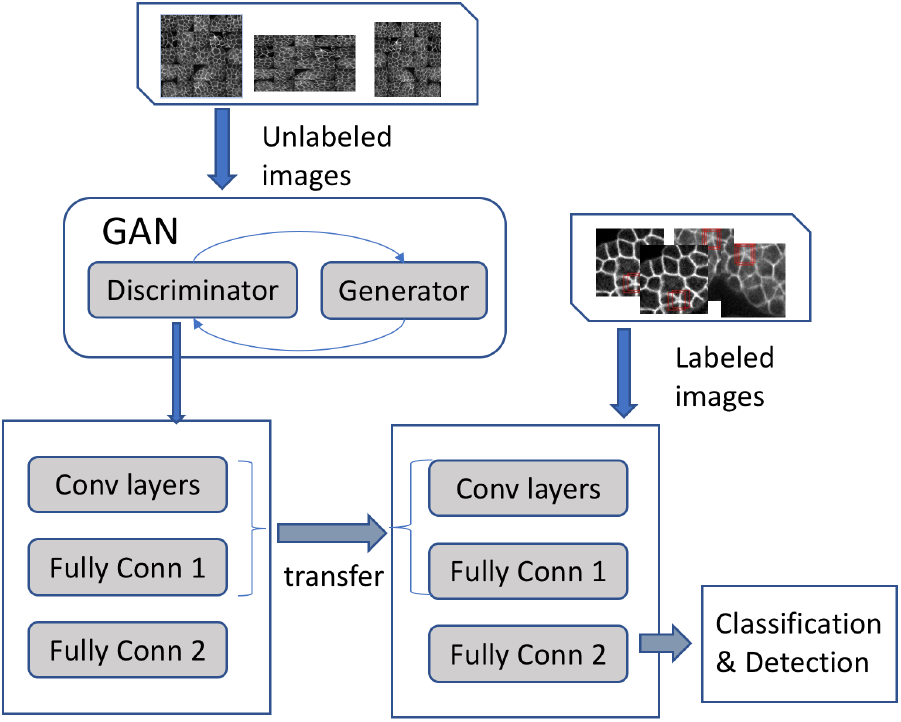
The overall workflow of GAN-based image classification scheme. There are two steps: common feature learning and target feature learning. An Alex-style network structure shown in Fig. 2 is used for both the GAN framework and the GAN-based classifier.

### 2.5 Data augmentation and hyperparameters

Because of the limited number of labelled images, we use several techniques to compensate for the potential problems associated with small training datasets for image classification and pattern detection. Specifically, we apply dropout during training after the first fully connected layer to eliminate the over-fit problem. Furthermore, we apply several data augmentation techniques to our dataset including randomly flipping the image vertically or horizontally and adjusting the brightness and the contrast of the image by a random percentage in a certain range. We use a learning rate of 10^−5^ and a batch size of 32 for the training of the network.

### 2.6 Computational platform

We implement our networks with tensorflow 1.7.1, a publicly available deep learning framework. More specifically, the convolutional network for classifying is built upon tensorflow’s Estimator API with a convolutional network as the customized model function. The generative adversarial network is implemented with tensorflow’s TFGAN framework with both the generator and discriminator customized. All experiments are performed on an Nvidia DGX server with four cutting-edge Nvidia Tesla V100 GPUs. Each Tesla V100 is equipped with 640 Tensor cores and a 16 GB memory.

## 3 Results

### 3.1 GAN-based classifier is better trained

When using a small training dataset, it is known that a conventional classifier can run into a data over-fitting problem quickly (Zeiler and Fergus, 2014). Compared with a neural network that directly trained on the small labelled dataset, a GAN-based network achieves better testing accuracy and, more importantly, demonstrates a more stable training process. To better understand how the GAN-based classifier works differently from the conventional classifier, we investigate the weights of filters on the first convolutional layer (e.g., the conv1 layer in Fig. 2). Figure 5 shows the weights of the filters in the first convolutional layer of both a conventional classifier and a GAN-based classifier that are trained for the 32 x 32 images.

**Fig. 5.**
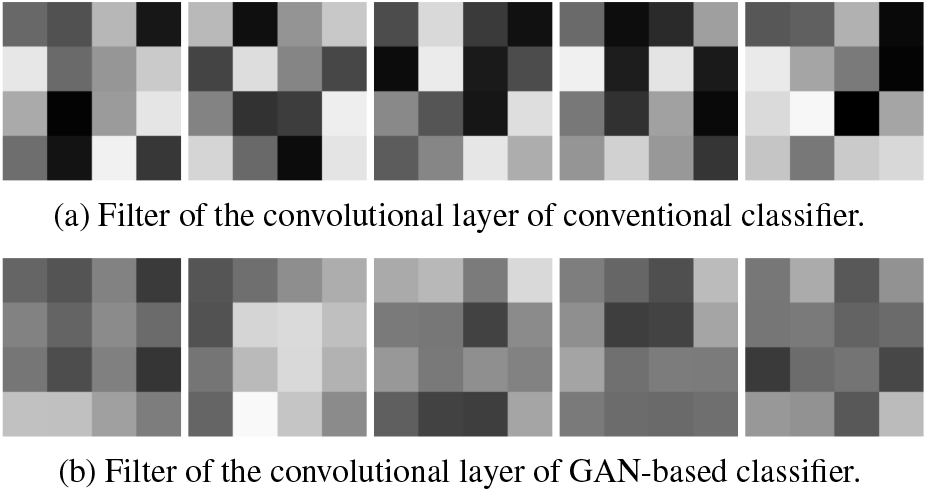
Visualization of the filters of a conventional classifier and a GAN-based classifier.

The weights of the GAN-based classifier (shown in Fig. 5(b)) are smoother than the weights of the conventional classifier (Fig. 5(a)). Quantitatively, we measured the standard deviations of both sets of weights and found that the STD of the GAN-based classifier is much smaller than that of the conventional classifier (0.192 vs. 0.285). The filter smoothness indicates that the GAN-based classifier is better trained.

We also analyzed the weights of the second-to-last fully connected (FC) layer (e.g., the fc1 layer in Fig. 2) of both the conventional classifier and the GAN-based classifier. There are 2048 x 256 parameters in the FC layer within the network for 32 x 32 images. The weights of the FC layer contain essential information on how the neural network handles the input images for the classification. Due to the complexity of a convolutional neural network, it is difficult to explicitly explain the role of these weights using a regular mapping function (for more information, see http://cs231n.github.io/understanding-cnn/). Here we reveal some differences between the conventional and GAN-based classifiers. We use the t-Distributed Stochastic Neighbor Embedding (t-SNE) method (Maaten and Hinton, 2008) to visualize the weights of these networks and explore their local similarity. The weight of the FC layer in the neural network for 32 x 32 images using t-SNE visualization is illustrated in Fig. 6.

**Fig. 6.**
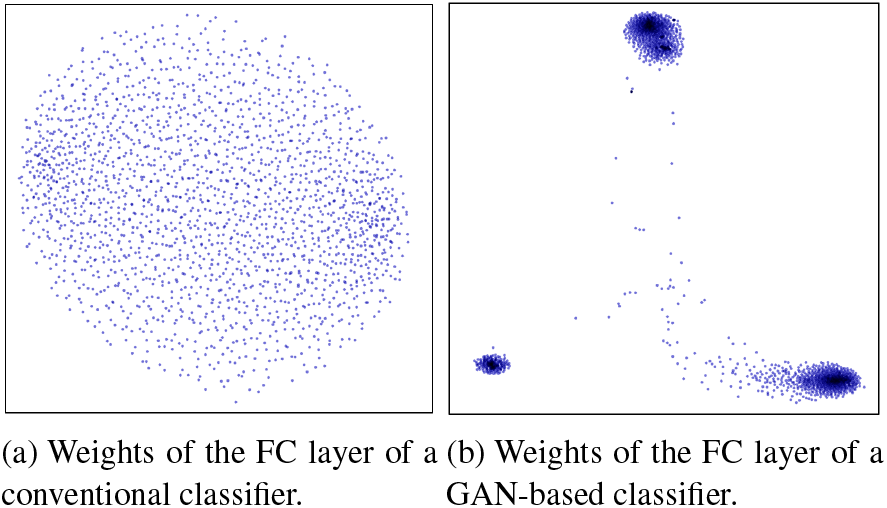
Visualization of the weights of FC layers in a conventional and a GAN-based classifier.

As shown in Fig. 6, the weights of the FC layer in the conventional classifier (illustrated in Fig. 6(a)) are more uniformly distributed, which means that the similarity of the individual weights is not significant. Compared with Fig. 6(a), Fig. 6(b) has three tight clusters, which infers that the GAN-based neural network is more sensitive to the subtle differences in specific structural features, such as the size of structural center and the contrast in edges.

We also compared visualizations of the output of each layer from both of the GAN-based model and the conventional model trained for the 32 x 32 images using the small dataset in Fig. 7. Because the values of the activations in each layer are not of the same scale and size, we first normalize all feature maps and then reshape them to a size of 32 x 32. In the visualizations of conv layer 2, we saw a clear pattern of a rosette in the GAN-based model, which cannot be found in the conventional classifier.

**Fig. 7.**
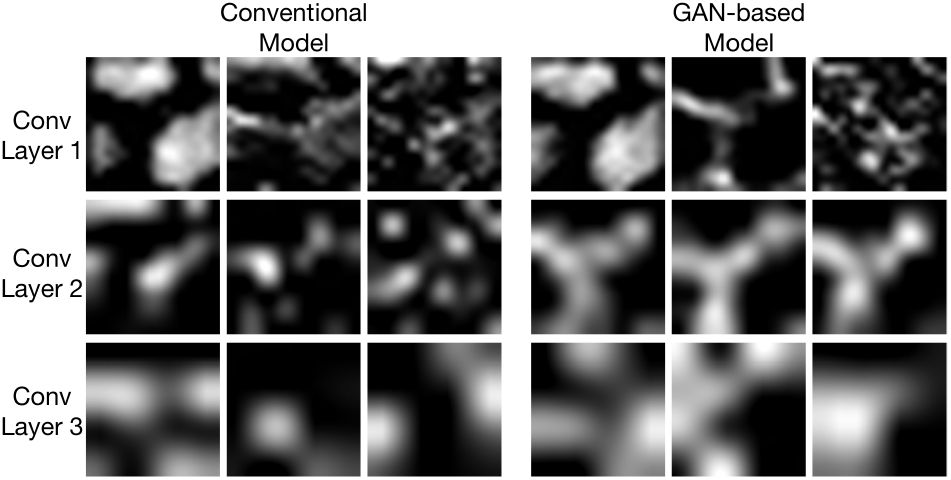
Visualizations of the output of each layer in GAN-based model vs. a conventional neural network with small training dataset.

From the visualizations we also found that the activation maps of the GAN-based model look brighter than those of the conventional model, especially in Layers 2 and 3. The magnitude of the activations (the values of the feature maps) is an reasonable indicator of how well a model is trained. These activations, working as feature detectors, with higher values are often more important to the classification task than those with lower values (Molchanov *et al.*, 2016; Li *et al.*, 2016). To measure the magnitude of activations quantitatively, we calculated the mean activations in each trained model using a small 32 x 32 dataset. We compared the mean activations at four different configurations of the training process: (1) random initialization without any training, (2) initialization with the parameters from the discriminator of the pretrained GAN but without any fine-tuning, (3) the trained conventional model, and (4) the trained GAN-based model (Fig. 8). We found that the mean activation value of each layer from a model without training is quite low. On the other hand, those values from a model with initialization by the pretrained GAN significantly increase, indicating a much better starting point for fine-tuning a labelled dataset. We then trained both models for 20k iterations over the 32 x 32 training dataset and found that the mean activations in the GAN-based model are larger than those in the conventional model.

**Fig. 8.**
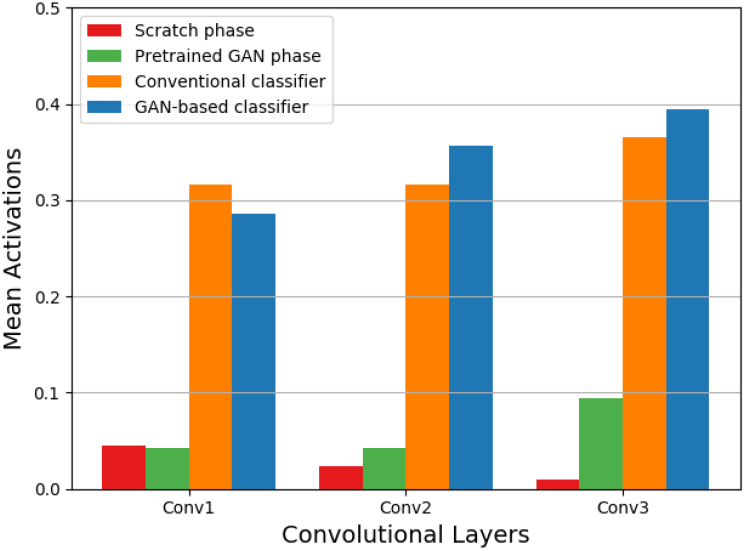
Mean activations of each convolutional layer inside different configurations with small training samples.

### 3.2 GAN-based classifier outperformance conventional classifier

#### 3.2.1 Experiments

As shown in the previous subsection, the GAN-based classifier is better trained than the conventional classifier using a small labelled dataset. This is significant for developmental biology since sometimes sizable labelled data is difficult to obtain. In this section, we first investigate the effect of the size of a training dataset on the performance of conventional and GAN-based classifiers. We then further evaluate the performance of the GAN-based classifier with training datasets of a different size.

Two sets of experiments were designed to investigate the effect of the size of a training dataset on the performance of conventional and GAN-based classifiers. We have 78 32 x 32 rosette images. Because rosette images appear less frequently in the observation dataset than non-rosette images, we selected three times more non-rosette images (195 images) from these unlabelled datasets. Because these *C. elegans* images are collected on three different dates, we selected the annotated images (20 rosette images and 20 non-rosette images, the ratio of rosette to non-rosette is 1:1) collected on one specific day as the validation dataset. All the images from the other two days were used as a training dataset (58 rosette images and 175 non-rosette images, the ratio of rosette to non-rosette is around 1:3).

In the first set of experiments, we used the entire training dataset (233 images) and test dataset (40 images). In the second set of experiments, we only used around 1/5 of the training datasets (12 rosette image and 40 non-rosette images) to train the neural networks. We adopted all of the data augmentation techniques mentioned in Section 2.5 in each experiment.

We adopted the F1 score as a measure of our network testing accuracy. An F1 score considers both the precision and the recall of the neural network outputs to compute the harmonic average of the precision and recall. An F1 score reaches its best value at 1 (perfect precision and recall) and worst at 0. It is a good measure for our study because the rosette and non-rosette images have a unbalanced distribution within our datasets.

#### 3.2.2 Performance

The model performances of the two sets of experiments are shown in Figure 9. The upper graphs (Fig. 9(a)) show the model performance with the entire training dataset, while the lower graphs (Fig. 9(b)) show the performance of the second set of experiments using 1/5 of training dataset (12 rosette and 40 non-rosette images).

**Fig. 9.**
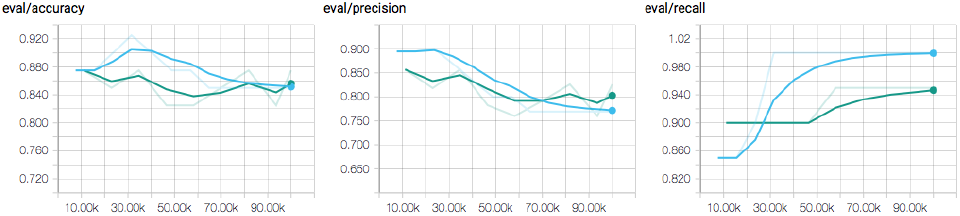
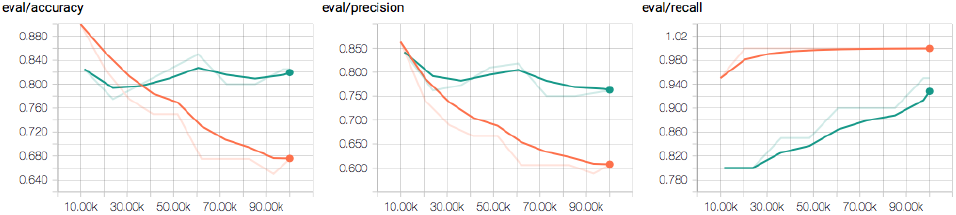
The comparison of model performance with the entire (upper graphs) and 1/5 (lower graphs) of the training dataset. The graphs on the left show the total accuracy of both rosette and non-rosette image prediction, while the graphs in the middle and on the right show the model prediction and recall rate of rosette data only.

As shown in the graphs on the left in Fig. 9(a), the GAN-based network and the conventional classifier perform the same with the entire dataset. The prediction accuracy over the 52 32 x 32 images is around 86%.

The left (accuracy) graph in Fig. 9(b) shows that the GAN-based classifier outperforms the conventional classifier to overcome the data underfitting problem. A decrease in the precision of conventional classifier prediction (shown in the middle graph of Fig. 9(b)) is the main reason for performance deterioration. It is also worth mentioning that the GAN-based classifier works pretty well using the small training datasets, with an accuracy of around 82%, which is comparable with that of the entire training dataset (shown in Fig. 9(a)).

### 3.3 GAN-based classifier sustains performance better with small data samples

We further investigated the impact of dataset sizes on the GAN-based classifier by conducting 11 sets of experiments using the whole (78 rosettes and 195 non-rosettes) or partial training datasets (100%, 90%, 80%, 60%, 50%, 40%, 30% 20%, 10% and 2%). In each of the experiments, we conducted 20 individual runs. Each used either the entire dataset or a randomly generated partial training dataset. The results are illustrated in Fig 10.

**Fig. 10.**
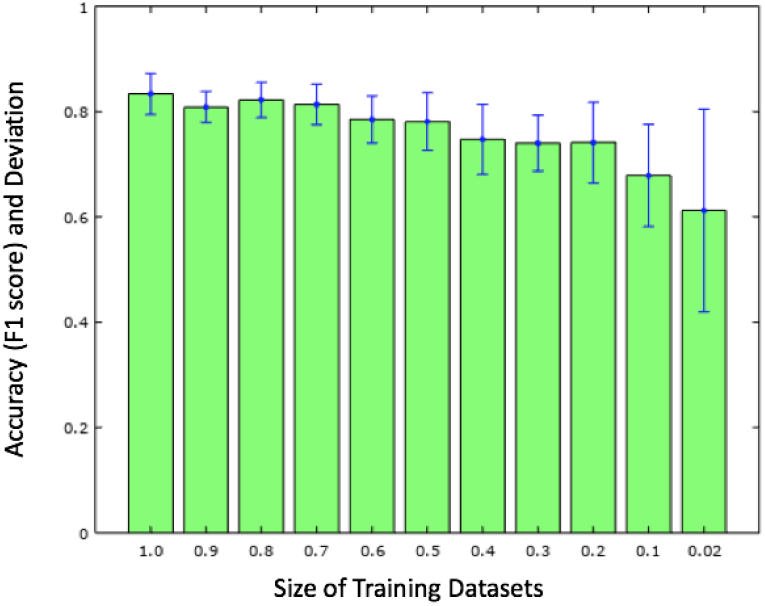
The accuracy and standard deviation of GAN-based classifier with datasets of different sizes.

As shown in Fig. 10, the average accuracy of the GAN classifier with the entire dataset is around 84% with a deviation of 4%. The model accuracy is around 80% (with a standard deviation of 4.5%) when we use more than 40% of the entire training set. The accuracy decreases to 76% and the standard deviation increases to 7% when only 20% of the training dataset is used (i.e., 12 rosette images and 40 non-rosette images). When the training dataset is too small (10% or less of the dataset), the accuracy drops much faster and the standard deviations become much larger. Further evaluations over the precision and recall of these experiments (when over 20% training datasets are used) reveal that the decrease in accuracy is mainly due to a decrease in precision; that is, when the training data is insufficient, the classifier identifies many rosette images, but some of the predicted images are non-rosettes when compared with the training labels. In a summary, Fig. 10 shows that the GAN-based classifier can deliver comparable accuracy (over 90% of the accuracy with the entire dataset) using around 1/5 of training datasets. These results indicate that significant time can be potentially saved in training data annotation and preparation.

### 3.4 Rosette detection

A well-trained GAN-based classifier (with a small 32 x 32 dataset) can be used for rosette detection inside large observation images. It can also be used for large image classification (such as an image with or without a rosette).

We use a sliding window (32 x 32) approach to identify and detect a 32 x 32 block of the 128 x 128 images. We adopt a stride of 4 pixels; therefore, there are 24 x 24 steps for each 128 x 128 image. At each step, if the model predicted a higher probability value than a predefined threshold (such as 0.9), we drew a red box around the current image block. We recorded all the probability output of each scan and generated a probability heatmap to illustrate the results of rosette identification and detection. Two examples of the results are shown in Fig. 11.

**Fig. 11.**
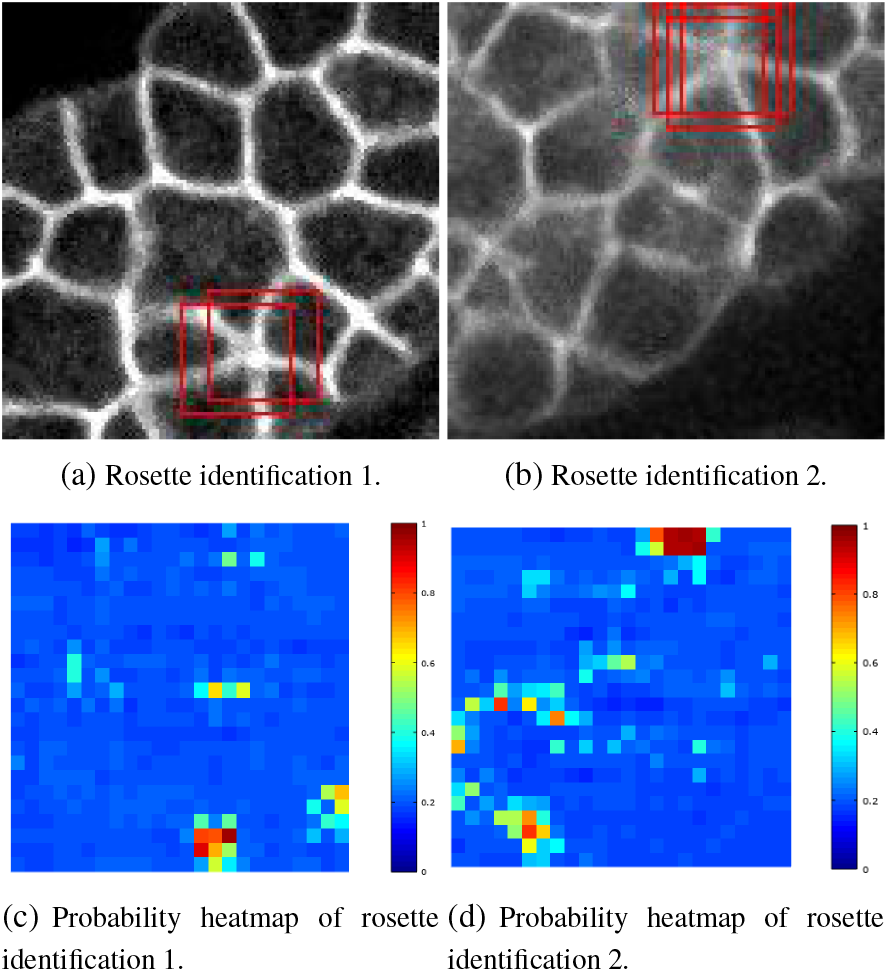
Example rosette images and associated probability heatmap for rosette detection.

### 3.5 Rosette dataset

We created an annotated dataset that includes around 400 128 x 128 images with explicitly marked 32 x 32 rosette structures. We further grouped the rosette images into two categories according to the probability value from the neural network. There are 85 images identified with a probability value greater than 0.9 and around 300 images with a probability value between 0.80 and 0.9. Examples of these images are illustrated in Fig. 12.

**Fig. 12.**
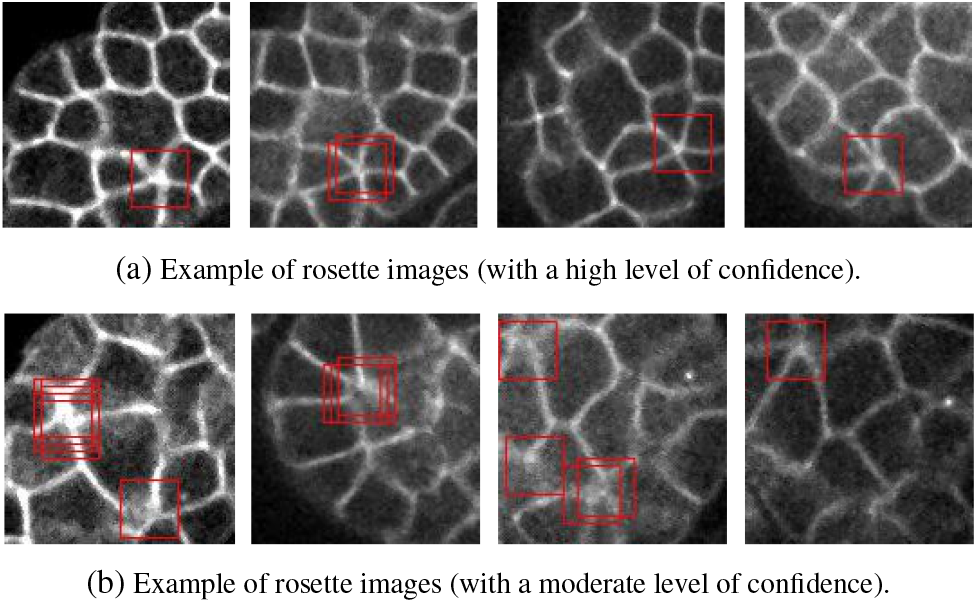
Rosette detection with high (upper graph) and medium probability (lower graph).

## 4 Conclusion

Compared with multi-institutional efforts for large-scale data exploration (Rajkomar *et al.*, 2018), establishing an adequate training dataset from a limited collection of microscopic images is a challenging but critical step to enabling deep-learning-based pattern identification. We have presented a GAN-based approach to efficiently classifying images with a particular cellular structure with relatively small unlabelled images and minimal annotated samples. By taking advantage of a competitive discrimination procedure with an unlabeled dataset, our GAN-based classifier can be better trained with small annotated training samples. Therefore, the the GAN-based classifier can outperform a conventional classifier (with same network structure) with a small annotated training dataset. Further quantitative measurements proved that the GAN-based classifier can sustain satisfactory performance even when 15 rosette and 36 non-rosette images were used. We think the methodology and concepts can be applied to other groups who are interested in using deep learning to identify and detect unique structures within microscopic data, such as flies, mice, and human brains.

## 5 Data and software availability

We created an annotated dataset that includes 128 x 128 images with explicitly marked 32 x 32 rosette structures. The dataset is available in dropbox (www.dropbox.com/sh/vlz3m2uzw73svts/AADHQpVGKNwnEskRF21oGJKTa?dl=0). These images can serve as a benchmark training dataset for further algorithm and application improvements. Related code and software utilities are located at https://github.com/daliwang/BioGAN.

## Funding

This study is partially supported by an NIH research project grants (R01GM097576). Research in the Bao lab is also supported by an NIH center grant to MSKCC (P30CA008748). This research used computing resources of the Oak Ridge Leadership Computing (OLCF) Facility and the Computing and Data Environment for Science (CADES) at the Oak Ridge National Laboratory, which is supported by the Office of Science of the U.S. Department of Energy under Contract No. DE-AC05-00OR22725.

